# Evaluating the toxicity of methanol, formaldehyde and formate on the growth fitness of *Yarrowia lipolytica*

**DOI:** 10.1101/2025.02.17.638576

**Authors:** Wei Kuang, Yuxiang Hong, Zimu Hu, Peng Xu

## Abstract

Liquid C1 feedstock, including methanol, formaldehyde and formic acid can be easily manufactured from greenhouse gases CO_2_ and CH_4_. *Yarrowia lipolytica*, isolated from marine ecosystem, is a non-conventional yeast which is capable of assimilating multiple range of hydrophobic compounds including alcohols, lipids, hydrocarbons, and volatile fatty acids *et al*. This yeast has been extensively engineered to produce lipids, carotenoids, squalene and other natural products. Herein we tested the growth fitness of *Y. lipolytica* when exposed to various one-carbon (C1) liquid feedstocks (methanol, formaldehyde, and formic acid). We firstly captured the cell growth kinetics with an automatic Cell Growth Analyzer and fitted the data with a Monod-type growth model. We obtained the inhibition constants of the three liquid C1s from the growth fitness dose-response curves. We found that the lagging phase duration is exponentially correlated with the toxicity of the tested C1 solvents. Notably, *Y. lipolytica* exhibited tolerance to methanol (IC_50_ 871 mM) and formic acid (IC_50_ 42.6 mM), but showed sensitivity to formaldehyde (IC_50_ 3.8 mM). We next chose fromate as the co-substrate to cultivate *Y. lipolytica*, confirmed that *Y. lipolytica* can upcycle formic acid to improve lipid yields and biomass in fed-batch culture. This work broadens *Y. lipolytica*’s substrate scope and facilitates sustainable and cost-efficient production of fuels and chemicals.

## 1. Introduction

Metabolic engineering is the targeted modification of cell metabolism to synthesize high-value products. These commonly-engineered cell factories are predominantly derived from industrial workhorse yeasts and bacteria, which are modified using genetic tools to incorporate functions not inherent to their native metabolic pathways [1]. The economic value added through this innovative process is derived from the difference between the cost of the raw materials and the market value of the final products. Currently, the primary substrates utilized by yeasts are fermentable sugars including glucose (C_6_H_12_O_6_), fructose, pentose or other complex carbohydrates, which naturally elevate the production costs in metabolic engineering. However, C1 feedstocks, such as methanol, formaldehyde, and formic acid, have garnered increasing attention in metabolic engineering community due to emerging technologies that enable their synthesis from simple and efficient process [2]. A notable example is the water-gas shift reaction, a well-established industrial process that converts carbon monoxide and water into carbon dioxide and hydrogen gas. This reaction not only efficiently produces hydrogen, which is a crucial feedstock for various downstream processes, but also facilitates the generation of various C1 feedstocks (methanol, formaldehyde and formate), when the released CO_2_ is electrochemically reduced [3]. In addition, greenhouse gas methane (CH_4_) or CO_2_ can also be selectively oxidized or reduced to form various C1 feedstocks in gas-liquid heterogenous phase with zeolite catalysts [4]. Electrochemical reduction and heterogenous catalysis reactions exemplify scalable and cost-effective processes to produce these simple liquid C1 feedstocks, which can subsequently be incorporated into cell metabolism through the diversified microbial carbon assimilation pathways [5]. Consequently, if non-sugar feedstock, such as liquid C1s can be assimilated as carbon sources to support cell growth and produce economically viable end-products, the cost of raw materials can be significantly reduced, thereby enhancing the competitiveness of the biomanufacturing and fermentation processes [6-8].

However attractive it might sound, one major issue regarding the efficient utilization of C1 feedstocks are their high toxicity to microorganisms. Many organisms, naturally, have the capability to capture and assimilate substrates like methanol and formate *via* native pathways [9]. Some methanotrophic bacteria, such as *Cupriavidus necator*, can directly uptake methane or formate due to the presence of Calvin-Benson cycle [10]. When encountered methanol in the surroundings, *C. necator* can first convert methanol to formaldehyde, where formaldehyde will later be incorporated into Calvin-Benson cycle and participate in the cellular metabolism. Other metabolic pathways, such as bacterial Ribulose Monophosphate cycle (RuMP cycle) which directly converts methanol or formate to glyceraldehyde 3-phosphate (G3P), alternate C1 assimilation route is the serine-glycine cycle which may couple the production of metabolic intermediate pyruvates and malates, also exists naturally in certain microbial systems [11]. However, due to the incomplete or inefficient C1 assimilation pathways, most yeasts or bacteria cannot process methanol, formate, or formaldehyde and incorporate them into their biomass [11]. Consequently, this largely hindered the development of utilization of C1 feedstock as the sole carbon sources to culture yeasts.

*Yarrowia lipolytica*, naturally found in oil refinery plant or marine environment, is a model organism for metabolic engineering due to its strong adaptivity to diverse conditions, ability to assimilate hydrophobic substrates and availability of facile genetic modification tools [12]. Wild type *Y. lipolytica* cannot utilize C1 carbons (methanol, formaldehyde and formate) as sole carbon sources [13]. This yeast has been extensively engineered to produce lipids [14], carotenoids [15], squalene [16], and other natural products [17, 18]. A number of studies have attempted to co-feed methanol or formate with glucose to facilitate green manufacturing of organic acids or lipids in *Y. lipolytica* [19, 20]. However, few studies have thoroughly examined how well *Y. lipolytica* can tolerate various liquid C1 carbons. For example, the Konzock’s research report [21] in 2021 showed the formic acid won’t have effects on *Y. lipolytica*’s growth until the concentration reaches 18.75 mM. However, in our high-throughput experiment, we found *Y. lipolytica*, without any pre-treatment, can tolerate up to 32 mM of formic acid (nearly 1.5 g/L) before an abrupt decline in growth fitness after this concentration limit. In this work, we used high throughput cell growth analyzer and automatically recorded *Y. lipolytica* growth under various concentrations of C1 feedstocks. By fitting the growth data into the Monod model, we were able to obtain the specific growth rate, and these rates can be regressed to a dose-response curve to derive the half inhibition concentration (IC_50_) of the tested C1 feedstock. We were also able to establish the exponential correlation between the extended lag phase and the toxicity of the C1 feedstock. Finally, we have co-fed formate with glucose to an engineered lipid strain, and we found the co-feeding strategy significantly improved cell growth and lipid production. The reported work investigates the toxicity of various C1 feedstocks, expands the substrate range of *Y. lipolytica* and may facilitate the upgrading of low-cost methanol or formate into value-added compounds through microbial conversion.

## 2. Materials and Methods

### 2.1 Yeast strains and Media

The *Po1g* strain was selected as the basal strain for subsequent C1 feedstock testing and lipid production. By integrating the NotI-linearized pYLXP’ and pYLXP’-ACC1-DGA1 [22] plasmids into the PBR322 docking platform of *Y. lipolytica* [23], we created the control strain Po1g-pYLXP’ free of leucine auxotrophic marker and the lipid production strain P01g-ACC1-DGA1. Yeast extract-peptone-dextrose (YPD) broth, consisting of yeast extract (10 g/L; OXOID), peptone (20 g/L; Sangon Biotech), and dextrose (20 g/L; Aladdin), was used as the base medium for growing the yeast strains. The plasmid pYLXP’-ACC1-DGA1 [22] was constructed previously and stored at −80 °C freezer in a medium containing 50% glycerol and 50% YPD broth in the lab. Under a biosafety cabinet, 3 ml of pre-prepared YPD solution was transferred into a 10-ml **plastic tube**; an aseptic inoculation ring was used to scrape some frozen yeast mixture and transfer it into the tube. Afterwards, the tubes were placed in a **shaking incubator** at 30°C and 250 **rpm** for 48 hours. This growth medium containing the viable yeast cells was then used as the inoculum.

### 2.2 Automatic Growth Analyzer and OD Measurements

Before the inoculation of Po1g-pYLXP’, the standards solutions of methanol (24.47 M, 5 M, 1 M; CAS 67-56-1), formate (25.97 M, 0.1 M, 0.05 M; CAS 64-18-16), formaldehyde (14.16 M, 0.05 M; CAS 50-00-0) were diluted with sterile water. After 2 days of growth, the OD600 of inoculum were in the range of 2.4 to 3.3 (**Thermo scientific Genesys 150**). The inoculum was further diluted using prewarmed YPD broth (30 °C) to achieve a final OD600 of approximately 0.1 into each **entry** of the sterile **48-deep well plates**, with pre-filled C1 liquid feedstock (methanol, formate, formaldehyde), to achieve the following concentration gradients: methanol (0, 1, 2, 4, 8, 16, 32, 64, 128, 256, 512, 768, 1024, 2048, 4096 mM), formaldehyde (0, 0.5, 1, 2, 4, 8 mM), and formate (0, 1, 2, 4, 8, 16, 32, 64 mM). These concentration gradients were determined with a few rounds of pre-experiments. The final volume of each well is 1mL. Then the plates were placed in a shaking incubator (30°C; 800 rpm) with Automatic Growth Recording function (MicroScreen, High-Throughput Microbial Growth Curve Analysis System, Gering Instrument Manufacturing (Tianjin) Co., Ltd., China), which monitored the OD600 of each well for every 10 minutes. Each concentration of C1 feedstock was tested in triplicate to minimize random variation and single-point anomalies. After 96-120 hours of incubation, all OD600 data points were pooled into the raw data spreadsheet for kinetic analysis.

### 2.3 Sigmoidal yeast growth fitting curve

After acquiring the data points, the software Excel (https://www.microsoft.com/en-us/microsoft-365/excel) was initially utilized to compute the expected values and standard errors for each concentration of the C1 feedstocks. Subsequently, the preprocessed data were imported into the software OriginPro (https://www.originlab.com). To model the OD as a function of time, a non-linear four-parameter dose-response growth curve was selected and depicted as follows:

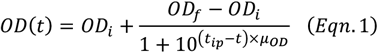

In this context, *OD* refers to *OD*_600_ value, *t* denotes time. *OD*_*i*_ and *OD*_*f*_ represent the lower and upper asymptotes of the fitting curve, corresponding to the initial and final OD values, respectively. *μ*_*OD*_ is the maximal specific growth rate *μ*_max_ associated with OD values, and *t*_*ip*_ is the time at the inflection point of the fitting curve. In practice, the initial OD measurement of each sample ranged from 0.22 to 0.32, with one outlier above 0.3, predominantly clustering between 0.23 and 0.25. This variation is primarily due to the time elapsed between the inoculation of each well and the reading delay of the optical measurement by the machine, as well as the manual handling involved, which could result in a deviation of 20 to 30 minutes (from OD value of 0.1) in the initial measurement.

### 2.4 Maximal specific growth rate determination

The maximal specific biomass accumulation rate of yeast (*μ*_max_; the reciprocal of time 1/h) provides insights into the growth fitness of a certain type of organism under a fixed set of conditions. The specific growth rate (*μ*) is defined as

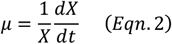

where *X* is the biomass concentration (g/L). According to the Monod equation that states

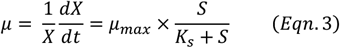

where S represents the limiting substrate concentration (g/L) and *K*_*s*_ denotes half saturation constant. Knowing the yield coefficient (*Y*_*X*/*S*_) is identified as

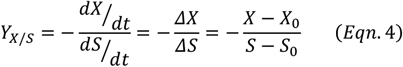

and so, converting it to get a function of 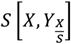

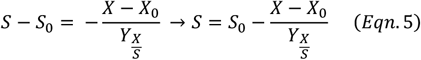

the Monod equation can be modified to

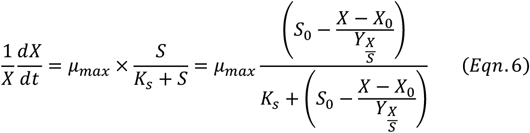

and the biomass accumulation rate (g/L/h) is of the final form [24]

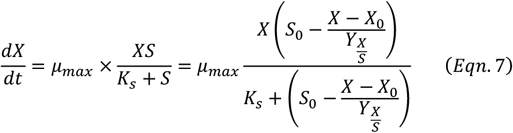

where *S*_0_ and *X*_0_ are the initial amount of substrate and biomass concentration, respectively.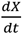 can be approximate to 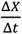, where Δ*t* is a fixed time interval (10 min) between each two measurements. The optimal density *OD*600 can be regarded as an effective parameter for biomass concentration, in general, the relationship between OD and biomass follows a linear relationship, *X* = *λ* × *OD*, where *X* denotes the biomass concentration (mg/L) and *OD* represents the dimensionless OD600 value. *λ* here denotes the conversion factor between biomass and OD value, which normally ranged **between 0.3 g/L and 0.5 g/L per OD unit**. The selection of OD in this study wouldn’t influence the tolerance analysis of C1 feedstock because the linear nature of two variables. Thus, an arbitrary conversion factor *λ* = 0.3 was chosen. As for the initial biomass concentration *X*_0_, even though the first OD measurement of each sample ranged from 0.22 to 0.32, considering the time elapse between the inoculation and the first measurement, we estimated the OD of culture right after the inoculation should be in the range from 0.15 to 0.2 (corresponding to 0.045 g/L to 0.06 g/L). Therefore, a reasonable *X*_0_ of 0.05 g/L was deliberately selected. After altering the biomass concentration to g/L, sample data are further processed in Matlab (http://www.mathworks.com/downloads/) to numerically solve these three parameters (*μ*_*max*_, *K*_*s*_, and *Y*_*X*/*S*_) by reversely fitting the OD∼time raw data into the above-mentioned Monod model using nonlinear least square regression.

### 2.5 Relative growth fitness (percentage of *μ*_*max*_), half time (*t*_50_), and lag time (*t*_20_, modeling

The time *t*_50_ denotes the amount of time the yeast culture takes to reach half-way of the maximal cell density. For each substrate, the *t*_50_ against C1 inhibitor concentration was plotted and modelled to find the best fit curve. The model [25] we used is

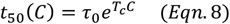

where *τ*_0_ denotes the time to reach half maximal cell density without any C1 inhibitors. The parameter *T*_c_ measures the **toxicity level** of the corresponding C1 substrate. The same fitting model was also applied to demonstrate the relation between **lag time** *t*_20_ and concentration, where *t*_20_ represents the time takes to reach 20% of the maximal cell density. *μ*_*max*_ stands for the maximal specific growth rate under certain amount of C1 inhibitor compared to the case without inhibition.

To plot the *μ* _*max*_ percentage against concentration, we used a sigmoidal type dose-response model [26] to showcase the maximal specific growth rate and the inhibitor concentration:

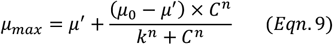

In this dose-response curve, *μ*^*′*^ is the maximal specific growth rate at high concentration of inhibitor and *μ*_0_ is the basic maximal specific growth rate without any inhibitors. *k* represents the IC_50_ value, which is the concentration of inhibitor when 50% of the maximal specific growth rate is achieved. *n* denotes the cooperation coefficient between yeast *Yarrowia lipolytica* and the corresponding C1 inhibitor. And C is the inhibitor concentration (mM).

### 2.6 Lipid production and analysis in the glucose media with formate co-feeding

The engineered strain ACC1-DGA1 was used as a control strain [27]. For lipid production, the yeast was cultivated under C/N ratio 80 media containing 40 g/L glucose, 1.1 g/L (NH_4_)_2_SO_4_, 1.7 g/L Yeast Nitrogen Base (YNB) and 0.74 g/L CSM-Leu (Sunrise, CA), in a total working volume of 30 ml per each flask system. As a comparison, two strains were supplemented with 20 mM (1.36 g/L) or 50 mM (3.4 g/L) formate. The cells were cultivated at 30 °C with shaking at 250 rpm. Samples were taken everyday up to 6 days. The lipid extraction and analysis follow the protocols reported previously [28].

## 3. Results and discussions

### 3.1 Failed adaptive lab evolution to acclimate *Y. lipolytica* to C1 feedstocks

Out of curiosity, in this work, we first endeavored to cultivate the yeast *Yarrowia lipolytica, Po1f strain*, to gradually acclimate and thrive in a culture of methanol, formaldehyde and formate. The Po1f strain was isolated from a culture containing 4.0 g/L YNB-AA+AS, 1 g/L yeast extract, and 10 g/L glucose after 48 hours. We next conducted an adaptive laboratory evolution process by transferring the strains to successively higher concentrations of methanol and formate (and lower concentrations of glucose) in the culture medium every 48 hours. Notably, except in the first few rounds where glucose was added to ensure a dense and viable strain could be harvested, in the subsequent adaptive steps, only methanol, formaldehyde and sodium formate were used as the sole carbon sources for the growth of the Po1f strain. We measured the optical density (OD) before each transfer to monitor the strain’s growth. However, on average, after few rounds for methanol and formic acid, first round for formaldehyde, the cells stop growing and the experiments ended unsuccessfully: we observed that the cells experienced a significantly extended lag phase without growing when methanol, formaldehyde or sodium formate were used as the primary carbon sources.

Following the *ex post* profiling, the researchers hypothesized that the extended lag time of the yeast *Y. lipolytica* after each transfer might be due to the toxicity of the tested C1 carbon sources. The lag phase is a complex preparatory period during which yeast or bacterial cells acclimate to a new environment [29]. Several factors influence the duration of this phase, including nutrient availability, environmental toxicity, and the history of the inoculated strains. This lag phase is crucial for yeast cells as it allows them to adjust their transcriptome and proteome before initiating cell division. According to some literature [30], cells may even reduce in number initially due to environmental challenges. In the previous trials, we speculated that 48-hour lag time was insufficient for the yeast cells to recover and propagate. Consequently, a new research direction was established to study the toxicity of C1 feedstocks and the lag period of *Yarrowia lipolytica* when inoculated in a co-culture of glucose and C1 feedstocks, which will help us determine how different amounts of C1 feedstocks (methanol, formate, and formaldehyde) influence the lag time and growth fitness of *Y. lipolytica*.

### 3.2 *Y. lipolytica* growth fitness with methanol as co-substrate

The growth curve analysis with methanol as a co-substrate highlights the remarkable tolerance of the yeast *Yarrowia lipolytica* to methanol. At a concentration of 512 mM (approximately 16.4 g/L), the proliferation of *Y. lipolytica* remains unaffected, showing no significant inhibitory effects. However, as the methanol concentration increases, inhibitory effects become evident, primarily through a prolonged lag phase. Repeated assays confirm that the *Y. lipolytica* can proliferate even at a maximum concentration of 2048 mM (65.6 g/L), with notable growth occurring after approximately 80 hours. Fig. 1 clearly shows that the variability in data points becomes more pronounced beyond 768 mM (approximately 24.6 g/L). This variation suggests that different yeast populations may exhibit distinct lag phases as they acclimate to the methanol-induced stress before entering the propagation phase [11]. Generally, the more toxic the environment, the longer it takes for yeast strains to adjust their cellular machinery (e.g., transcriptomes and proteomes), as illustrated by the growth trajectories in the figure.

**Fig 1.**
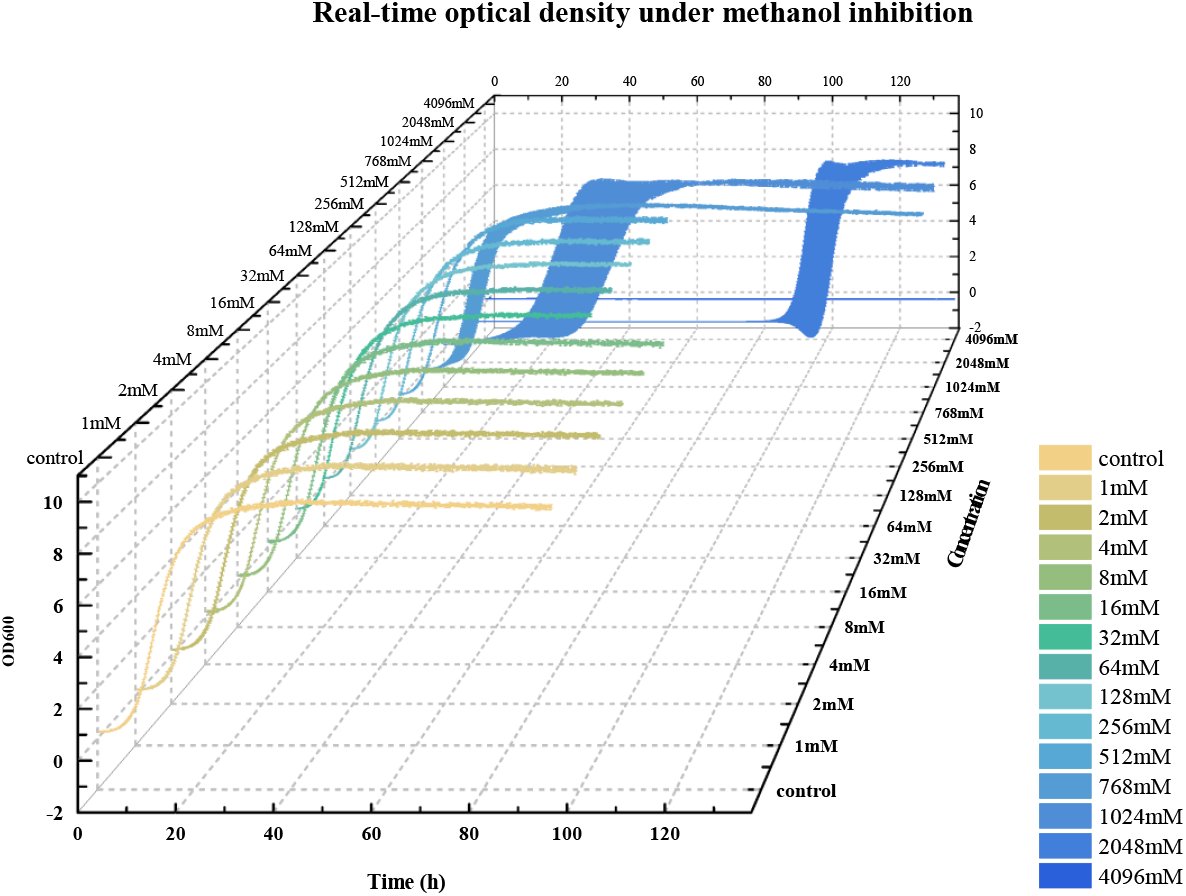
*Y. lipolytica* growth curve with various concentration of methanol.

Interestingly, methanol appears to be less toxic to *Y. lipolytica* than previously believed. The strain demonstrates robust growth, particularly at methanol concentrations below 768 mM (24.6 g/L). The tolerance to MeOH is possibly due to the oleagineity nature which may help this yeast maintain its membrane integrity in the presence of methanol [31]. This finding suggests that *Y. lipolytica* may possess a highly effective detoxification mechanism, allowing it to mitigate the toxic effects of methanol and maintain normal cellular respiration, with negligible impact observed at concentrations below 15 g/L.

### 3.2 *Y. lipolytica* growth fitness with formaldehyde as co-substrate

Formaldehyde, in contrast to methanol, exhibits a profound inhibitory effect on the proliferation of *Y. lipolytica*, as depicted in **Fig. 2**. The yeast strain demonstrates a markedly lower tolerance to formaldehyde, with growth inhibition becoming apparent at concentrations as low as 0.5 mM. The maximum concentration at which the yeast can maintain some level of growth is 4 mM (approximately 0.12 g/L), beyond which growth is severely inhibited, if not completely halted, particularly noticeable at 8 mM where no significant growth is observed over the 80-hour period. This results suggest that the bottleneck of efficient formaldehyde/methanol utilization is the lack of a strong formaldehyde sink pathway (for example formaldehyde dehydrogenase or xylulose monophosphate pathway XuMP pathway) [11] in *Y. lipolytica*

**Fig 2.**
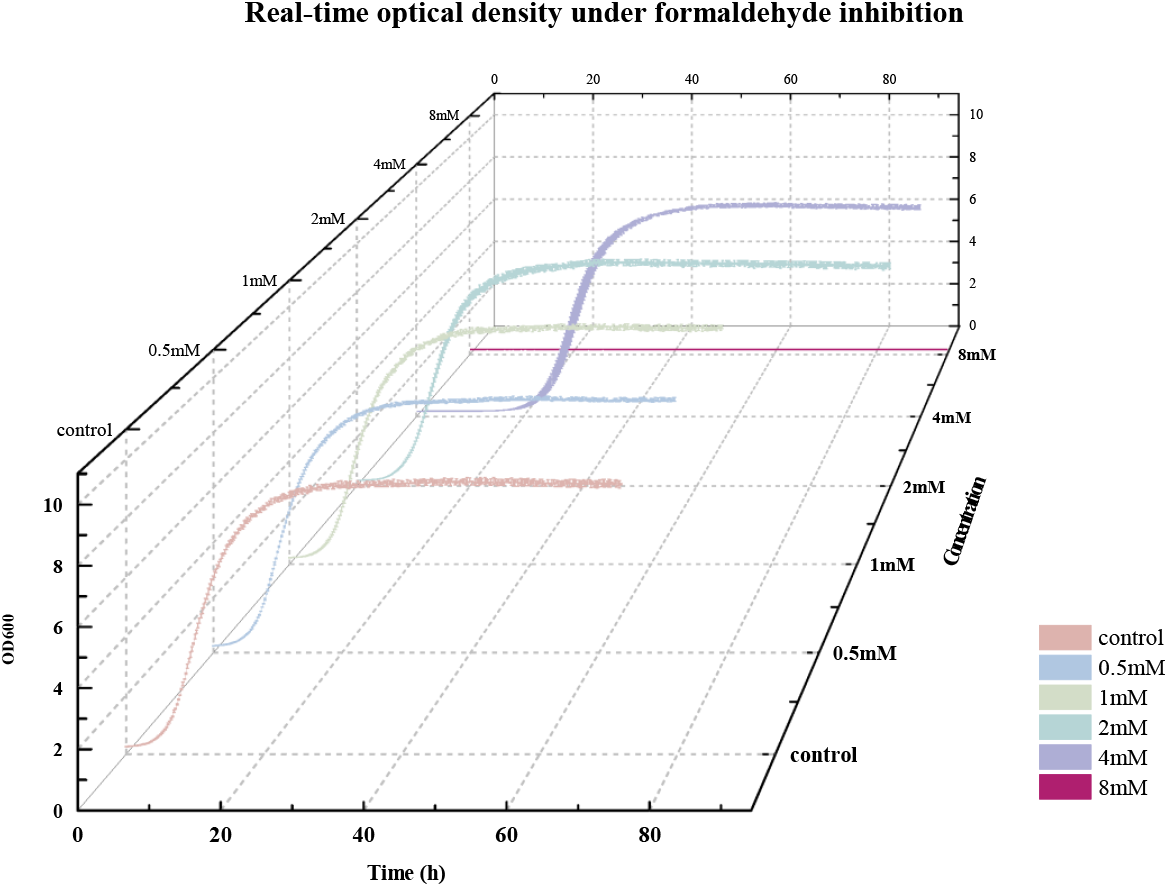
*Y. lipolytica* growth curve with various concentration of formaldehyde.

This stark contrast in tolerance compared to methanol can be attributed to the highly reactive nature of formaldehyde. As a potent electrophile, formaldehyde readily reacts with nucleophilic sites in cellular macromolecules such as proteins, nucleic acids, and lipids [32]. This reactivity disrupts essential cellular processes, including DNA replication, transcription, and protein function, leading to cellular stress and, ultimately, growth inhibition. The toxic effects are compounded by formaldehyde’s ability to form cross-links within and between these macromolecules, further impeding cellular function and viability [33].

Moreover, the delayed growth at lower concentrations of formaldehyde, as observed in Fig. 2, suggests that *Y. lipolytica* attempts to adapt to the presence of formaldehyde through various stress response mechanisms, albeit with limited success at higher concentrations. For example, when low concentration of formaldehyde (lower than 4mM) was applied to the media, the dwindled (not completely ceased) growth might result from an intrinsic anti-oxidative defense mechanism in *Y. lipolytica*, however, after the threshold amount, the mechanism invalidates, and the yeast cannot maintain its growth fitness. The inability to proliferate at concentrations above 4 mM underscores the severity of formaldehyde’s cytotoxicity, highlighting its potential to cause substantial cellular damage even at low concentrations. This makes formaldehyde a particularly challenging substrate for *Y. lipolytica* and suggests that any metabolic engineering efforts to utilize or tolerate formaldehyde would require significant modification to the yeast’s natural detoxification pathways.

### 3.4 *Y. lipolytica* growth fitness with formate as co-substrate

Simultaneous tolerance testing for formate using the yeast strain *Y. lipolytica* has yielded noteworthy results [19], demonstrating that *Y. lipolytica* can tolerate up to 32 mM (approximately 1.5 g/L) of formic acid with minimal inhibitory effects. This tolerance suggests a robust adaptation mechanism in *Y. lipolytica*, potentially linked to the activity of endogenous formate dehydrogenase enzymes [34], which catalyze the oxidation of formate to carbon dioxide, thus mitigating the toxic effects of formic acid. However, as formic acid concentrations rise above 32 mM, the yeast’s growth exhibits a steep decline. At 48 mM, growth is severely inhibited, and at 64 mM, the yeast strain experiences complete cessation of growth, as observed in **Fig. 3**. The tolerance to formate may be ascribed to the six endogenous formate dehydrogenases [35] which can oxidatively decarboxylate formate to generate CO_2_ and NADH in *Y. lipolytica*.

**Fig 3.**
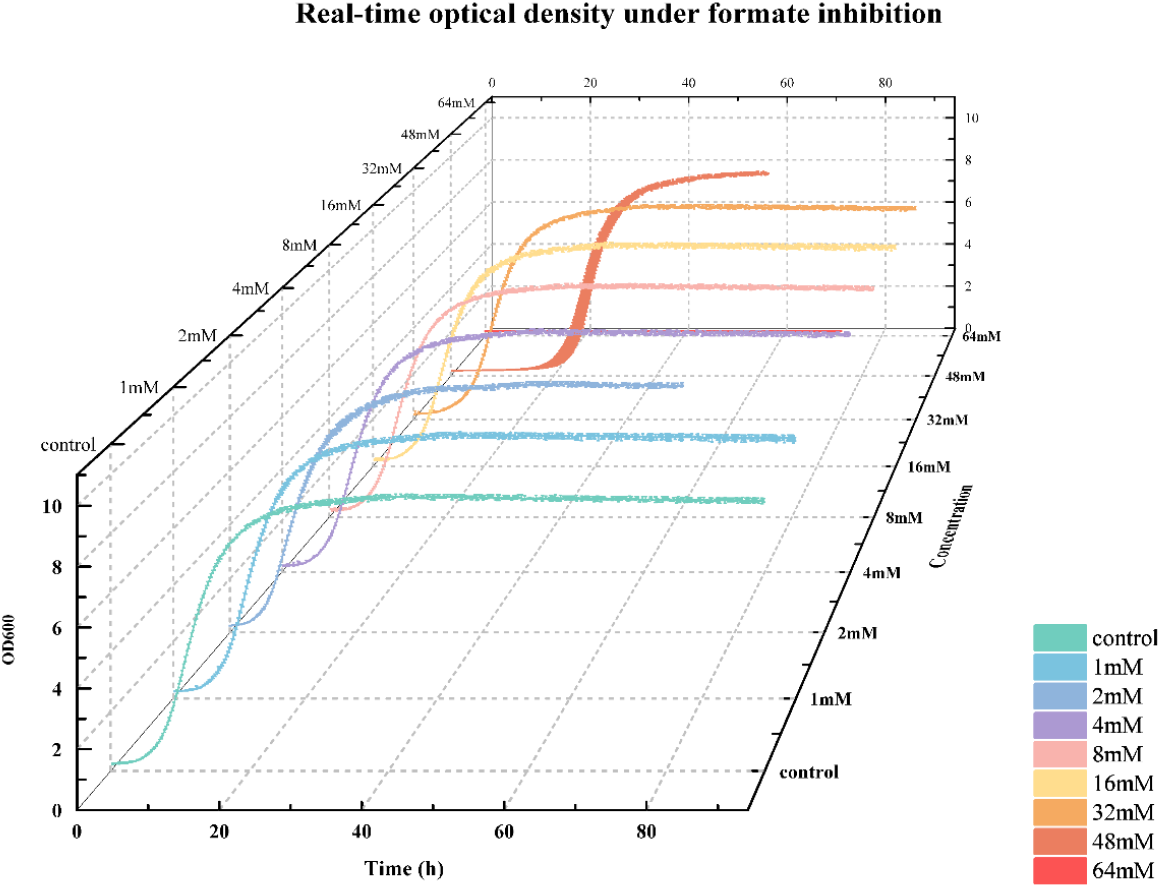
*Y. lipolytica* growth curve with various concentration of formate.

This abrupt transition from normal growth to complete inhibition likely indicates the presence of a critical threshold, beyond which the cellular detoxification pathways, including those involving formate dehydrogenase, are saturated. Intracellular accumulation of formate may lead to the disruption of cellular pH homeostasis, membrane integrity and interference with key metabolic processes [36]. While *Y. lipolytica* possesses endogenous formate dehydrogenase to degrade formate, its tolerance toward formate has a well-defined threshold (20∼48 mM), suggesting that any biotechnological applications involving formate as a substrate would need to carefully control formate levels to avoid surpassing this critical inhibitory threshold.

### 3.5 Evaluating the inhibition constant (IC_50_, of C1 feedstock on *Y. lipolytica*

**Fig. 4** illustrates the specific growth rates of *Y. lipolytica* under various concentrations of the C1 inhibitors methanol, formaldehyde, and formic acid, with the corresponding curve-fitting data summarized in **Table 1**. The dose-response curves and their fitted parameters reveal important insights into the inhibitory effects of these compounds. Each inhibitor displays a characteristic sigmoidal dose-response, with the IC_50_ value (denoted as “*k*” in Table 1) serving as a critical indicator of toxicity. The IC_50_ values distinctly rank the inhibitors in terms of toxicity, with formaldehyde showing the highest toxicity (*k* = 3.8 mM), followed by formic acid (*k* = 42.63 mM), and methanol demonstrating the least toxicity (*k* = 871.37 mM). This trend aligns well with the expected chemical reactivity and cellular toxicity of these compounds.

**Table 1.**
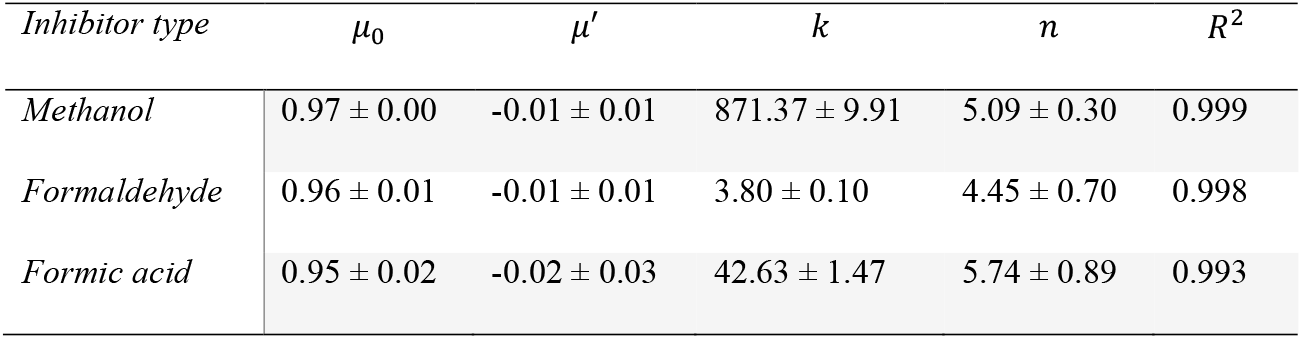
Fitting the growth data into the dose-response curve (*Eqn. 9*) to estimate the kinetic parameters of *Y. lipolytica* grown on methanol, formaldehyde, and formic acid.

| Inhibitor type | $\mu_0$ | $\mu'$ | $k$ | $n$ | $R^2$ |
| --- | --- | --- | --- | --- | --- |
| Methanol | $0.97 \pm 0.00$ | $-0.01 \pm 0.01$ | $871.37 \pm 9.91$ | $5.09 \pm 0.30$ | 0.999 |
| Formaldehyde | $0.96 \pm 0.01$ | $-0.01 \pm 0.01$ | $3.80 \pm 0.10$ | $4.45 \pm 0.70$ | 0.998 |
| Formic acid | $0.95 \pm 0.02$ | $-0.02 \pm 0.03$ | $42.63 \pm 1.47$ | $5.74 \pm 0.89$ | 0.993 |

**Fig 4.**
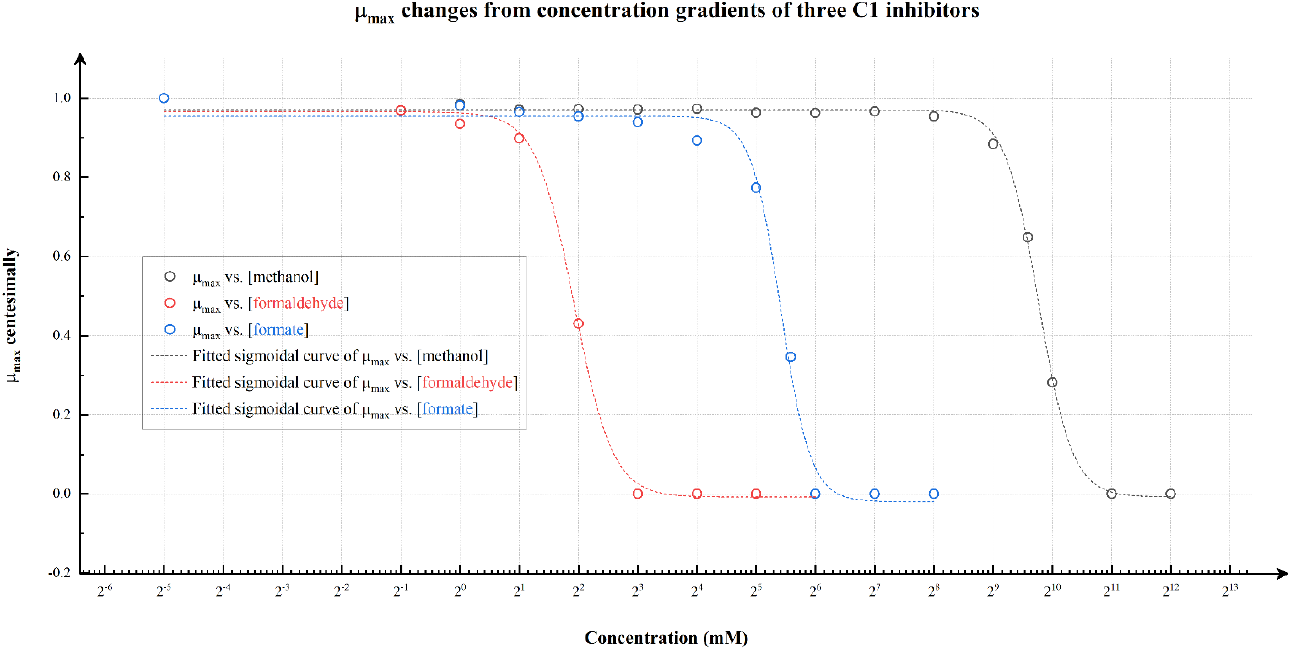
Dose response of relative growth fitness under various concentration of C1 feedstock

Furthermore, the cooperativity coefficient “*n*”, which reflects the interaction between the inhibitor and the yeast cells, varies among the inhibitors. Formic acid has the highest cooperativity coefficient (*n* = 5.74), suggesting a more complex or synergistic inhibitory interaction at higher concentrations, whereas methanol and formaldehyde exhibit slightly lower cooperativity values (*n* = 5.09 and 4.45, respectively). Although the exact biochemical basis for these differences in cooperativity is not fully understood, it may be related to the distinct modes of action of each inhibitor on cellular processes.

Notably, the maximal specific growth rate (*μ*’) approaches zero at high inhibitor concentrations, indicating complete growth inhibition due to the overwhelming toxicity. The control conditions for each inhibitor type yield nearly identical specific growth rates, confirming that the observed differences in growth dynamics are solely attributable to the presence of the inhibitors. The dose-response curves and kinetic parameters underscore the potent inhibitory effects of formaldehyde, the moderate inhibition by formic acid, and the relatively mild impact of methanol on *Y. lipolytica*, thus providing a clear hierarchy of toxicity among these C1 inhibitors.

### 3.6 Exponential correlation between toxicity and lagging phase duration

Using the collected data, the time required for each sample to reach half of its terminal optical density (OD), denoted as *t*_50_, was computationally calculated. For accuracy, OD measurements were replicated three times under identical conditions to mitigate the effects of single-point variations. **Fig. 5** illustrates that the relationship between inhibitor concentration and *t*_50_ follows a robust exponential correlation for all three inhibitors—methanol, formaldehyde, and formic acid—with adjusted R^2^ values exceeding 0.9. This strong correlation indicates that as the concentration of each inhibitor increases, the time required for *Y. lipolytica* to adapt to the adverse environment and reach half of its maximal population correspondingly increases.

**Fig 5.**
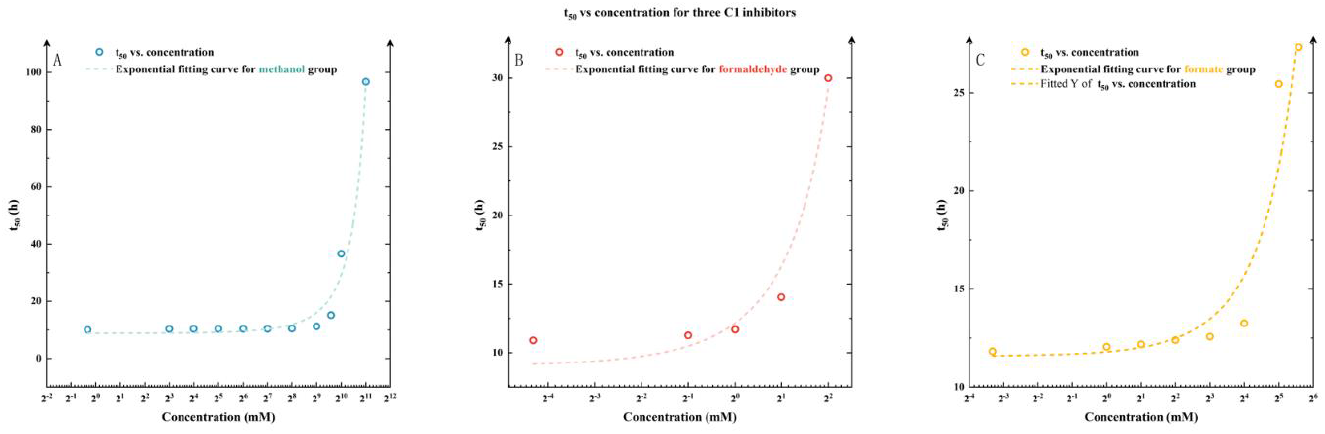
Exponential correlation between toxicity and lagging phase duration for three C1 inhibitors: methanol (A), formaldehyde (B), and formic acid (C).

According to the specific values presented in **Table 2**, despite a relatively large standard error (exceeding 0.72 hours) attributed to inherent growth inconsistencies among different yeast samples, it is evident that the time required for *Y. lipolytica* to reach half-maximal cell density in the absence of any C1 inhibitors (denoted as *τ*_0_) ranges from approximately 9 to 11 hours. This variation likely reflects the intrinsic variability in the growth kinetics of the yeast under optimal conditions.

**Table 2.** Fitting *t*_50_ as a function of the strength of the methanol, formaldehyde, and formic acid inhibitor.

| Inhibitor type | $\tau_0$ | $T_c$ | $R^2$ |
| --- | --- | --- | --- |
| Methanol | $8.809 \pm 0.998$ | $0.001 \pm 0.000$ | 0.98 |
| Formaldehyde | $9.051 \pm 0.914$ | $0.294 \pm 0.032$ | 0.96 |
| Formic acid | $11.574 \pm 0.720$ | $0.019 \pm 0.002$ | 0.91 |

The parameter *T*_*c*_ in the exponential model represents the toxicity level of each C1 inhibitor on *Y. lipolytica*. Methanol, with the lowest *T*_*c*_ value (0.001), exhibits relatively mild toxicity, meaning that gradual increases in methanol concentration only slightly affect the *t*_50_ of yeast growth. On the other hand, formaldehyde shows pronounced cellular toxicity, as evidenced by its significantly higher *T*_*c*_ value (0.294), which is about 25 times greater than that of methanol. This higher toxicity is further highlighted by the fact that *Y. lipolytica* can only tolerate formaldehyde concentrations up to 4 mM; beyond this threshold, growth is entirely suppressed.

In conclusion, the analysis of *t*_50_ and the associated parameters reveals distinct differences in how *Y. lipolytica* responds to these C1 inhibitors. Methanol is the least toxic, allowing the yeast to maintain relatively stable growth even as concentrations increase. Conversely, formaldehyde poses a significant challenge to yeast survival, sharply increasing the lag time and ultimately inhibiting growth entirely at higher concentrations. These findings underscore the critical importance of understanding the specific interactions between *Y. lipolytica* and each C1 inhibitor, particularly when optimizing industrial processes involving these substrates.

### 3.7 Cofeeding formate with glucose to improve lipid yield

Fig. 6 illustrates the impact of formate co-feeding on lipid production in *Y. lipolytica* when glucose was used as primary carbon source. For the control strain (Fig. 6A), the yeast efficiently utilizes glucose, as indicated by the sharp decline in glucose levels over time. The biomass increases steadily, reaching a peak around 50 hours. Concurrently, lipid production also increases, peaking at approximately 1.5 g/L. This baseline establishes that the yeast can effectively convert glucose into biomass and lipids under standard growth conditions.

**Fig 6.**
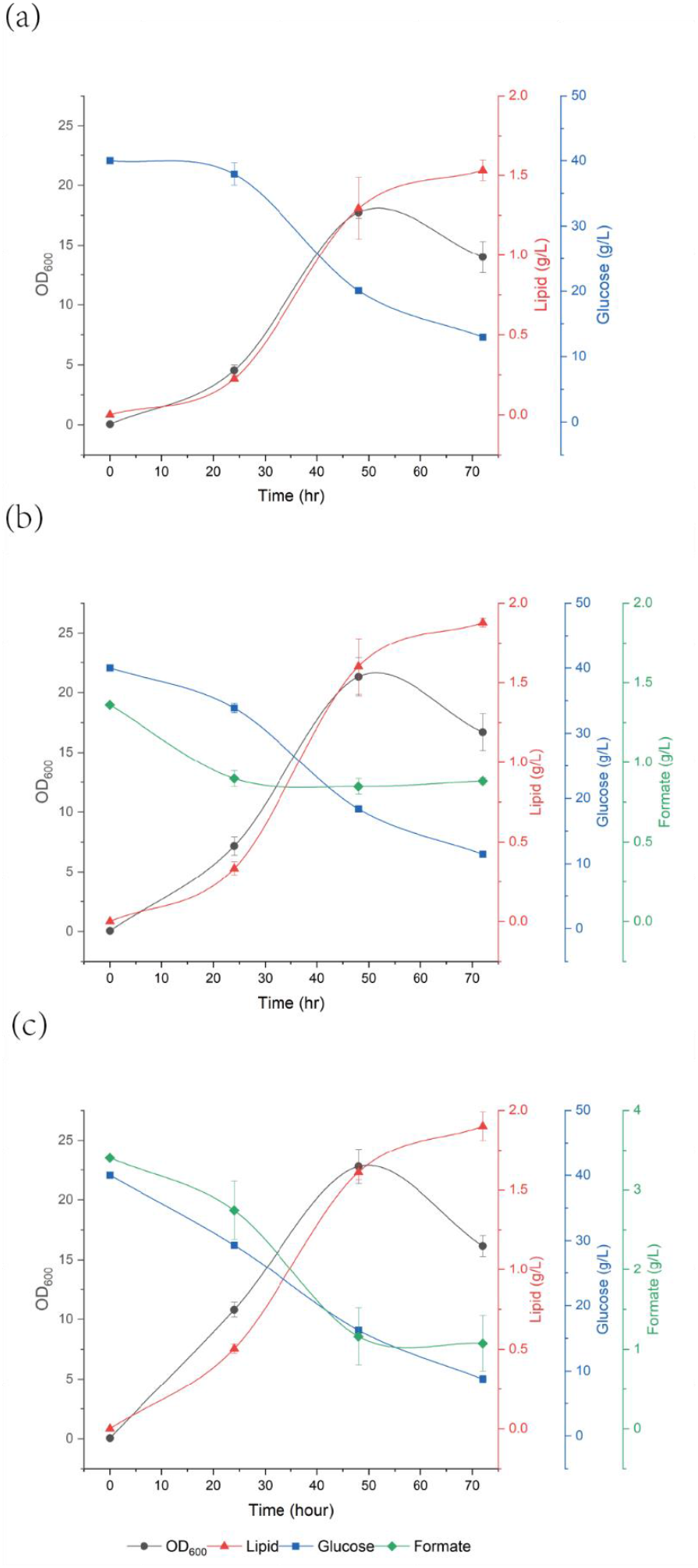
Lipid production of *Y. lipolytica* with glucose as carbon source co-fed with formate. A: 0 mM formate; B: 20 mM formate; and C: 50 mM formate.

When *Y. lipolytica* is co-fed with 20 mM formate (Fig. 6B), there is a noticeable enhancement in lipid production, which peaks at 2.0 g/L, a 33% increase from the control condition. This suggests that co-feeding with formate positively influences lipid biosynthesis. Interestingly, although formate is being consumed (Fig. 6B), the glucose consumption rate appears slightly delayed compared to the control. Furthermore, the overall biomass accumulation remains comparable, indicating that formate serves as an additional carbon source or NADH source, which supplements the glucose pathway facilitating lipid synthesis. Besides, through the well-documented POM (pyruvate-oxaloacetate-malate) cycle, NADH can be easily converted to NADPH which can promote lipid accumulation in *Y. lipolytica*.

When the formate concentration is increased to 50 mM (Fig. 6C), the yeast grows at slightly reduced rate compared to the lower formate concentration (Fig. 6B), possibly due to that the formate concentration is surpassing the IC_50_ value (42.6 mM) for *Y. lipolytica*, causing some inhibitory effects. Lipid production remains robust, peaking around 2.0 g/L, similar to the 20 mM condition. The glucose consumption rate is further delayed, and formate utilization appears to slow down as well, suggesting a possible inhibitory effect as formate concentration nears the IC50 threshold. Nonetheless, the data imply that even at inhibitory concentrations, formate co-feeding does not drastically hinder growth or lipid production, and in fact, still provides a benefit to enhance lipid production.

Overall, these results highlight the potential of co-feeding formate to improve lipid production in *Y. lipolytica*. The lipid yields have been increased from 0.056 g-lipid/g-glucose (no formate addition), to 0.066 g-lipid/g-glucose (20mM formate as co-substrate), to 0.061 g-lipid/g-glucose (50mM formate as co-substrate). While formate at lower concentrations (20 mM) significantly boosts lipid biosynthesis without compromising growth, even higher concentrations (50 mM) near the inhibitory threshold can sustain enhanced lipid production. Based on the performance of glucose uptake rates, which drops from 0.75g/hr (no formate), to 0.64 g/hr (20mM formate), to 0.54 g/hr (50mM formate), we can see a clear decline in the consumption of glucose for the yeast to grow; this further demonstrates the beneficial effect of formate in promoting cell growth. Additionally, after examining the lipid content percentages to cell biomass, we also can observe an increasing trend from 14% (no formate co-substrate), to 22% (20mM formate co-substrate), and to 23% (50mM formate co-substrate), also signifying that co-feeding of formate benefits lipid accumulation. This suggests that strategic co-feeding of formate could be a viable approach to optimize lipid production in industrial applications, making the process more efficient and cost-effective by utilizing an inexpensive and readily available carbon source.

## 4. Discussion and conclusions

The use of cell factories to produce valuable bioproducts has seen widespread application in recent years, extending from pharmaceuticals to consumer goods. This metabolic process, which transforms low-cost materials into high-value products, is revolutionizing the field, prompting the integration of novel genetic tools and innovative chemical reactor designs. One promising direction is the reduction of substrate costs, typically dominated by glucose, by substituting it with more economical and simpler compounds. Notably, some research teams have genetically modified *Y. lipolytica* to enable methanol assimilation from barely detectable level to a significant level of 1.1 g/L of biomass within 72 hours, by introducing and improving cellular RuMP/XuMP regeneration, formate dehydrogenation, and serine pathways [31]. From S. Zhang et al’ report, they even managed to enhance the methanol assimilation of *Y. lipolytica* to a level of 3.4 g/L with the synergistic carbon sources of methanol and xylose [13]. From our original high throughput analysis of the wild type of *Y. lipolytica* (without any genetic modifications), we have complemented the jigsaw by quantifying the maximal tolerancing behaviors of wild type *Y. lipolytia* toward various C1 feedstocks, which sets the foundation for future modifications. The simplest organic compounds—methanol and its derivatives, formic acid and formaldehyde—are ideal candidates for this approach due to their minimal structural complexity and low production costs. These three C1 substrates are byproducts of various industrial processes, potentially offer a novel path for reducing feedstock cost.

In this study, we investigated the tolerance of the *Y. lipolytica* to three inhibitors: methanol, formaldehyde, and formic acid, each known for its high cellular toxicity, particularly formic acid and formaldehyde. Using standard YPD media as the primary source for cell mass accumulation, we found that *Y. lipolytica* could tolerate up to 871.4 mM (approximately 27.9 g/L) methanol, indicating its significant potential as a primary carbon source for cellular growth. For formaldehyde and formate, the tolerance levels were 42.6 mM (approximately 1.96 g/L) and 3.8 mM (approximately 0.11 g/L), respectively, underscoring their substantial cellular toxicity, especially in the case of formaldehyde. Interestingly, under and up to 32 mM of formate, the inhibitory effects were negligible, comparable to growth without inhibitors. However, beyond 64 mM of formate, *Y. lipolytica* exhibited no growth, suggesting the presence of a unique cellular mechanism that protects the cells from formate-induced stress below 32 mM— a mechanism delicate enough to collapse immediately after this threshold is surpassed. Future research could aim to elucidate and validate this hypothesis.

*Y. lipolytica*, a model organism in metabolic engineering, demonstrates versatility in numerous applications for biomanufacturing and environmental remediation. A promising strategy to further reduce production costs involves identifying alternative carbon source for cell growth. Utilizing inexpensive and widely available C1 compounds as primary substrates for sustaining cellular activities and accumulating cell mass could significantly advance the pursuit of a more sustainable and environmentally friendly world.

## Acknowledgement

This work was supported by the National Natural Science Foundation of China (no. 22378083) and the Guangdong Provincial Key Laboratory of Materials and Technologies of Energy Conversion (MATEC, GR2300014).

